# Density-dependent spatial patterns and habitat associations of a dominant desert ant

**DOI:** 10.1101/2025.06.10.658907

**Authors:** Ana L. Cao, Fernando A. Milesi, Rodrigo G. Pol, Lucía Vullo, M. Florencia Miretti, Gabriela I. Pirk, Javier Lopez de Casenave

## Abstract

The abundance and distribution of ant colonies can reflect the relative importance of different ecological processes such as abiotic limitations and biological interactions varying with time, space and scale. We fitted a series of incrementally complex spatial models for point patterns in one and two dimensions to examine how the habitat associations and spatial arrangement of *Pheidole bergi* colonies change among scenarios of natural and anthropogenic variations in environmental conditions in the central Monte desert. At the habitat scale, *P. bergi* preferred the protected open woodland (algarrobal), with strong and persistent density variations at scales of tens of meters along most dirt roads crossing it. The 40% reduction in active colonies during the 2019–2025 drought was inversely density-dependent, increasing aggregation. Instead, colonies were 40–50% lower but more homogeneously arranged on roads crossing habitats with extensive natural or anthropic limiting factors (sand dunes and livestock-grazed algarrobal). Colony density within the protected algarrobal also showed >50% reduction between the wet 2001 and the dry 2019, with their distribution always influenced by the two-phase mosaic structure typical of arid lands. This preference for inter-patch borders and open areas rather than the interior of woody vegetated patches probably explains why the intensive anthropic impacts associated with dirt roads had no notable effect on *P. bergi* density. Beyond that spatial dependence, significative distance-related pairwise interactions between colonies shifted from slightly attractive in 2001, resulting in clumped distributions, towards strongly repulsive by 2019, favoring more random and overdispersed spatial patterns, contrary to the expected correlation between intraspecific competition and colony density. *Pheidole bergi* seems flexible enough to establish colonies in different habitats and microhabitats although with distribution and population dynamics still shaped by natural and anthropic, direct and indirect environmental constraints such as soil texture, vegetation cover or resource availability affected by multi-year droughts. The inversed density-dependent thinning and shifting relevance of inter-colony interactions with climate fluctuations observed at different spatial scales highlight the complex interplay between environmental constraints and intraspecific dynamics in the population ecology of *P. bergi*, an abundant, omnivorous, aggresive and probably ecologically relevant ant species in the Monte desert.

## Introduction

The abundance and spatial distribution of ant colonies are influenced by a range of biotic and abiotic ecological factors and can serve as indicators of underlying processes (Hölldobler and Wilson 1990). Their changes over space, time and scale respond to variations and shifts in those processes and can elucidate the relative importance of concurrent environmental drivers. For example, intraspecific competition, which tends to increase with higher population density, is usually associated with regular colony distributions that minimize overlap in foraging areas of neighboring colonies (Bernstein and Gobbel 1979; Levings and Traniello 1981; MacKay 1991). Conversely, clumped distributions are more probable in unsaturated habitats and may result from differential colony survival induced by spatial heterogeneity in environmental factors such as vegetation cover, microclimatic conditions, altitude, soil type or food availability (e.g., Bernstein and Gobbel 1979; Nicolai 2005; Pol 2008; Milesi et al. MS). After some time, clumped patterns resulting from expansion during favorable reproductive periods or as legacy of initial habitat selection by founding queens may become more regular due to density-dependent mortality during harsher periods (Wiernasz and Cole 1995; Blanco-Moreno et al. 2014). And while coarse-scale aggregation may be expected from abiotic environmental limitations, negative interactions between neighboring colonies may reinforce regular distributions at fine-grained scales within suitable areas (Crist and Wiens 1996; Sundaram et al. 2022). Spatial patterns can not only reflect natural changes in environmental conditions, such as unusual multi-year severe droughts in deserts that affect both ant densities and their spatial distributions, but also responses to environmental modifications by anthropogenic factors (Crist and Wiens 1996; Sanders and Gordon 2004; Gibb et al. 2019; Sundaram et al. 2022).

*Pheidole* is the most diverse genus of ants, comprising approximately 1500 species distributed all over the world (except for Antarctica and some islands; Wilson 2003), and particularly abundant in deserts (MacKay 1991; Johnson 2001). In the Monte desert of Argentina they dominate the seed-carrying ant assemblages (Miretti et al. 2025), with *Pheidole bergi* as the most abundant and ubiquitous species in the central Monte (Milesi and Lopez de Casenave 2004; Claver et al. 2014; Miretti et al. 2025) and other arid and semi-arid areas in Argentina, Brazil, Paraguay and Uruguay (Kusnezov 1951; Wilson 2003; Wild 2007; Janicki et al. 2016; Guénard et al. 2017; Barrientos et al. 2024). It has an omnivorous, broad diet composed mainly of insects and other arthropods, supplemented with seeds and other plant materials (Pirk et al. 2009). Despite its recognized abundance, wide distribution, agresiveness towards other ants and tolerance to perturbations, little is known about its natural history and ecology (but see Wilson 2003; Pirk et al. 2009; Hierro et al. 2023). One such aspect is their association with habitat features and the spatial distribution of its colonies, and how they respond to temporal changes and perturbations that can affect the density of its populations in arid environments, such as prolonged droughts or anthropic impacts.

In the Monte desert, *P. bergi* confronts significant environmental heterogeneity at multiple spatial scales (Bertiller et al. 2009; Bisigato et al. 2009; Milesi et al. 2019). At landscape and habitat scales, geomorphological and edaphic factors influence vegetation structure, shaping habitats such as the “algarrobal”, an open woodland occurring on the deep soils of rolling planes, or strips of sand dune vegetation associated with coarse soils on elevated ridges (Lopez de Casenave 2001; Bisigato et al. 2009). Soil texture affects the abundance and distribution of many ant species (e.g., Kirkham and Fisser 1972; Johnson 1992, 2000; Boulton et al. 2005). At smaller scales, vegetation exhibits the typical two-phase mosaic structure of arid and semiarid landscapes (Aguiar and Sala 1999; Tongway and Ludwig 2005), i.e., shrub- or tree-dominated patches interspersed among herbaceous vegetation or bare soil, that strongly influences water and nutrients distribution, soil seed banks, micro-climatic conditions, and plant-animal interactions (Marone et al. 2004; Milesi et al. 2008, 2019; Bertiller et al. 2009; Bisigato et al. 2009; Andrade 2016). Anthropogenic disturbances further alter spatial heterogeneity at different scales: local perturbations such as site clearing and dirt roads modify micro environmental conditions, soil compaction and resources available for ants (Pirk et al. 2004), while extensive livestock grazing reduces plant richness and herbaceous cover, particularly grasses, expanding areas of bare soil (Milesi et al. 2002; Villagra et al. 2009; Pol et al. 2014; Andrade 2016). Although livestock grazing can have neutral or positive effects on some ant species (Bestelmeyer and Wiens 2001; Nash et al. 2004; Radnan and Eldridge 2018) it usually has negative impacts on ants (Nash et al. 2001; Goosey et al. 2019), including *P. bergi* (Calcaterra et al. 2010; Claver et al. 2014). As in most dry ecosystems, *P. bergi* populations must also cope with temporal heterogeneity dominated by infrequent and highly variable precipitation events (Noy-Meir 1973; Collins et al. 2014; Whitford and Duval 2020). Although seasonal distribution of rain is rather predictable in the Monte desert, the number and size of significant pulses and annual amounts are not, with series of consecutive dry years challenging productivity (Pol at al. 2010). In fact, a drastic reduction of perennial herbaceous cover has been recorded in the Ñacuñán Biosphere Reserve (Mendoza, Argentina) over recent decades (perennial grasses reduced their horizontal cover >75% since 2002; Milesi et al., unpublished data), with a particular sharp decline along the multi-year drought settled since 2018 (Miretti et al. 2024; Vullo et al. 2024).

In this study, we examined the abundance, habitat associations and spatial arrangement of *P. bergi* colonies under natural and anthropogenic variations in environmental conditions in the central Monte desert. At the habitat (macro) scale, we expected clustered spatial patterns in association with spatial variation in critical environmental variables such as soil type, water availability and correlated biotic factors that could affect food availability (e.g.: Johnson 1992, 1998; Nicolai 2005). At similar weather conditions, the degree of aggregation should be lower in milder habitats such as the main protected habitat in the Ñacuñán Biosphere Reserve (algarrobal) than in harsher environments such as similar areas under extensive anthropogenic exploitation (livestock grazing) or in habitats that naturally pose defying abiotic restrictions (the coarse and highly permeable soils in sand dunes), where suboptimal sectors are unable to sustain viable colonies (Bernstein and Gobbel 1979). Instead, the degree of aggregation in adequate habitats should decrease after harsh times (e.g, at the end of a multi-year drought), when colony density decreases faster in crowdier patches through intraspecific competition (density-dependent thinning; Sanders and Gordon 2004; Sundaram et al. 2022). At the patch (micro) scale we expected the distribution of *P. bergi* colonies to be strongly influenced by the vegetation mosaic, as reported for other desert ants (Carlson and Gentry 1973; Bestelmeyer and Schooley 1999; Nicolai 2005), but also for regular spatial patterns resulting from distance-dependent repulsion between neighbouring colonies (the intensity of foraging territory defense should increase with nest proximity for individually foraging ants, such as *P. bergi,* as opposed to those that forage following trunk-trails; Warburg and Steinberger 1997). Again, we expected the degree of regularity of those patterns to increase during the limiting conditions of a multi-year drought (2019) compared to more benign periods (2001), reflecting an increase in intercolony competition in dry years. To contrast the spatial configurations through those spatiotemporal scenarios at different scales and disentangle the potential underlying processes, we fitted and tested a series of incrementally complex spatial models for point patterns in one and two dimensions.

## Methods

### Study area

The study was conducted in the Biosphere Reserve of Ñacuñán (34°03’S, 67°55’W), located in the central Monte desert, Mendoza Province, Argentina. The region has a dry, highly seasonal climate, with warm and rainy summers and cold and dry winters. The mean (± SD) annual temperature is 16.8 ± 2.1°C, and the mean annual rainfall is 348.4 ± 123.4 mm (1972–2020). More than three quarters of the rainfall occurs in spring and summer (October–March), with annual accumulation strongly influenced by large summer rainfall events (>10 mm; Pol et al. 2010). Annual precipitation in 2018–2023 was approximately half of that recorded in 2000-2001, which in turn was part of a period of 5-6 years of above-average precipitation (Miretti 2022).

The main habitat of the reserve is the algarrobal, an open woodland of *Neltuma flexuosa* and *Geoffroea decorticans* trees scattered within a matrix of tall shrubs (mostly *Larrea divaricata*, but also *Condalia microphylla, Atamisquea emarginata* and *Atriplex lampa*), low shrubs (*Lycium* spp.*, Mulguraea aspera*, *Troncosoa seriphioides*), and perennial grasses (*Leptochloa crinita, Pappophorum* spp., *Sporobolus cryptandrus, Aristida* spp., *Digitaria californica, Setaria* spp., *Jarava ichu*). Forb cover (*Chenopodium papulosum*, *Glandularia mendocina*, *Parthenium hysterophorus*, *Phacelia artemisioides*, *Sphaeralcea miniata*) is highly variable among seasons and years. About one third of the algarrobal consist of open patches of variable size (up to several meters) without perennial vegetation (Marone and Horno 1997; Andrade 2016; Milesi et al. 2019). Other edaphic habitats within the reserve include northwest-southeast oriented sand dunes, stabilized by a similar woody shrub matrix (*L. divaricata*, *Ximenia americana*) with scattered *N. flexuosa* trees and some exclusive herbaceous species (*Panicum urvilleanum*, *Solanum euacanthum*, *Hyalis argentea*, *Gomphrena martiana*).

Livestock grazing, the main economic activity in this region, has been excluded from the reserve since 1972 (Pol et al. 2014). Surrounding cattle ranches share a similar floristic composition but exhibit lower herbaceous cover and smaller shrub and grass patches, resulting in larger areas of bare soil (Milesi et al. 2002; Andrade 2016). Perennial grasses are particularly affected by grazing, showing significant reductions in cover and seed abundance (Pol et al. 2014; Marone and Pol 2021). Both the reserve and nearby cattle ranches are crossed by dirt roads 6–9 m wide, with minimal rural traffic. Woody vegetation along these roads is periodically removed with heavy machinery. The altered soil surface is more compacted and exposed to sun, wind and rain than the adjacent woodland (Pirk et al. 2004).

### Sampling design

We assessed the abundance and spatial arrangement of *Pheidole bergi* colonies at different spatial scales across multiple years (2001, 2019, 2021, 2025). Sampling was conducted along dirt roads and within permanent grids in the Ñacuñán Reserve and a nearby cattle ranch. We always searched and mapped active colonies (i.e., with presence of workers) during February, at the peak of their external activity (Pirk et al. 2009). *Pheidole bergi* ants are small (major workers: 5.5–6.5 mm; minor: 3.2–4.5 mm; Bruch 1916; Wilson 2003) and many of their nests are rather inconspicuous (with a single, usually semicircular, small entrance 2.8–4.2 cm; Wilson 2003; personal observation), so we searched for colonies in at least pairs of trained people, twice during hours of their maximum foraging activity (6 to 3 h before solar midday). No field marks persisted between sampling periods.

At the macro scale (tens to hundreds of meters), to compare the same habitat (algarrobal) along a multi-year drought, in 2019 and 2025 we searched and mapped with a 100-m measuring tape (in one dimension; precision: 10 cm) all active colonies of *P. bergi* along two 1500 m segments (∼7 m wide) of different dirt roads (D and F) within the Reserve. Following the same protocol, to compare habitats under the same weather conditions, in 2021 we searched and mapped all colonies along three 500 m segments (6 m wide) of dirt roads located in each of three different habitats, two protected from domestic grazing within the Reserve (algarrobal and sand dunes) and one in an adjacent cattle ranch (grazed algarrobal).

At the micro scale (meters), in 2001 (wet period) and 2019 (dry period) we followed a similar search procedure to map (in two dimensions; precision: 1 m^2^) all active colonies of *P. bergi* within two permanent 50×50 m grids (J and V) established in the protected algarrobal, ∼80 m apart and >50 m from the nearest dirt road. To facilitate and ensure a thorough census of colonies, each grid was divided in 5×5 m squares with temporal marks. The limits between patches of woody vegetation taller than 1 m and open patches (inter-patch borders hereafter) were drawn on a gridded map of each area (except for V in 2001) and then forced to the same spatial grain (pixel: 1 m^2^). The distance to the nearest inter-patch border was calculated for each pixel center, with positive distances for pixels located in open patches and negative values for pixels within woody vegetation patches. Distances to vegetation for V in 2001 were assumed equal to those measured in 2019 so those particular results should be taken with more caution (linear correlation between distances in J in 2001 and 2019: r = 0.67).

### Spatial data analyses

The spatial distribution of *P. bergi* colonies was analyzed using spatial statistics for point patterns in one (roads, macro scale) and two (grids, micro scale) dimensions, implemented in several functions within the *spatstat* family of packages (version 3.3-2, Baddeley et al. 2016) in the R platform (R Core Team 2025; version 4.5.0). Full R code for data processing, exploration and analyses, including partial and additional results, is available at DOI:10.5281/zenodo.15521695 (in Spanish).

#### Colony distribution along dirt roads (macro scale)

We first explored spatial patterns by testing the null hypothesis of a Completely Spatial Random (CSR) processes, i.e., that the observed location of colonies along roads are outcomes of Poisson spatial processes with spatially-invariant intensity (the expected number of colonies per spatial unit). We tested for CSR with (1) a Kolmogorov-Smirnov goodness-of-fit test evaluating the maximum deviation of the observed cumulative distribution function of coordinates of colonies against the expected function under the CSR null hypothesis (Baddeley et al. 2016), and (2) second-order global functions (Ripley’s K and pair correlation) which are sensible to spatial autocorrelation resulting from a spatially heterogeneous variable that affects the intensity of the process (e.g., an important soil characteristic varying along the censused road segments) or from positive/negative distance-dependent interactions among colonies in a non-Poisson spatial process (or “true” autocorrelation; see Dale and Fortin 2005; Baddeley et al. 2016). Ripley’s K function performs better as a basis for statistical inference while the pair correlation function, being non-cumulative, is less ambiguous and easier to interpret (Baddeley et al. 2016). The empirical one-dimensional version of the K function (K^L^) estimates the average number of points (colonies) in 2*r* length segments centered at each focal point, standardized by the intensity so it can be compared among point patterns (Baddeley et al. 2016). Edge effects were geometrically corrected with a one-dimensional analog of the standard isotropic correction in the bidimensional case (Ang et al. 2011; Baddeley et al. 2016). To avoid artifacts from excessive edge correction, *r* values did not exceed a quarter of the segment length (Fortin 2020): *r* ≤ 350 m for 1500 m segments and *r* ≤ 120 m for 500 m segments. We used a centered version of the K function with zero as the expected value for a CSR process at all *r* distances, positive values indicating clustering and negative values, repulsion or overdispersion. The linear pair correlation function (g^L^) estimates the ratio between the average number of pairs of points separated by *r* and the expected number under a CSR process with the same intensity (the “instant” version or first derivative of the K function; Baddeley et al. 2016). For a CSR the expected value is 1, with higher values indicating aggregation (i.e., more pairs of points observed at *r* distance than expected by chance), and lower values, segregation. Confidence envelopes for both functions were obtained by Monte Carlo simulations with 199 iterations of CSR processes with the same intensity as the observed point pattern. The empirically estimated 95% critical envelopes at each *r* were used as local hypothesis tests only after a global test of Maximum Absolute Deviation suggested the rejection of a CSR model (Baddeley et al. 2014).

When the CSR hypotheses were rejected, we explored the nature of the detected autocorrelation among colonies with the inhomogeneous versions of the K and pair correlation functions. These modified functions control for the autocorrelation derived from variations in the expected number of colonies along the roads by estimating the intensity function with a non-parametric Gaussian kernel smoothing of the point pattern data. To test the null hypothesis that the spatial process is still Poisson (i.e., there are no “true” distance-related interactions among colonies), assuming the estimated intensity function, pointwise envelopes and global Maximum Absolute Deviation tests were estimated as above. As it has been shown that they may result in conservative tests (Baddeley et al. 2017), we also considered a Balanced Independent Two-Stage Monte Carlo test of goodness-of-fit for composite null hypotheses (Baddeley et al. 2017) with 1560 iterations (39 iterations of the first stage, each with 39+1 iterations for the second nested stage). No further increases in model complexity were necessary at this scale.

#### Colony distribution in the algarrobal (micro scale)

For two-dimensional point pattern, we analyzed spatial autocorrelation with Ripley’s L (a transformation of Ripley’s *K*: *L* = sqrt(*K*/π)) and pair correlation functions. The L function averages the number of points lying within circles of radius *r* centered on each of them, with values above expected CSR process suggesting aggregation, and values below, overdispersion or repulsion. The pair correlation function *g(r)* is similar but considering points separated exactly by each distance *r* (in its empirical version, averaging the number of points lying on thin rings of *r* radius centered at each point in the pattern); values above 1 suggest aggregation, and below 1, repulsion. A Ripley’s isotropic edge correction was applied to both functions: the point counts were weighted by the reciprocal of the proportion of the sampling circle’s circumference lying within the observation area, with *r* ≤ 12 m (∼1/4 grid sides) to control for excessive edge effects (Fortin and Dale 2005; Baddeley et al. 2016). Confidence envelopes for *L(r)* and *g(r)* were obtained by Monte Carlo simulations with 399 iterations of CSR processes with the same intensity as the observed point pattern, with the empirically estimated 95% intervals at each *r* used as point-wise hypotheses tests after a global test of Maximum Absolute Deviation suggested rejecting the null hypotheses (Baddeley et al. 2014).

We first explored the relationship between *P. bergi* colonies and woody vegetation with Kolmogorov-Smirnov goodness-of-fit tests, comparing the distributions of distances to inter-patch borders of pixels with colonies and those available in each of the grids. We evaluated habitat selection at this scale with Monte Carlo tests comparing the position (mean) and dispersion (standard deviation) of the distributions of distances between pixels with colonies and their nearest inter-patch borders against the distributions of those statistics obtained from 1999 random samples of the same size from the available distances in each grid (a CSR null model). Then, after several steps of data exploration (see available document with partial and additional results at DOI:10.5281/zenodo.15521695), we fitted Gibbs point process models of different complexity to the observed bidimensional spatial patterns of colonies, assuming the two grids as replicates of the same spatial process and considering Year as a fixed design factor (with two levels: 2001 and 2019) affecting one or more model parameters. Gibbs models explicitly postulate distance-dependent attractive (resulting in clustering) or inhibitive (repulsion or overdispersion) pairwise interactions between points, reducing to Poisson models (independent points) when interactions are nil. We modeled the log of the conditional intensity (the probability of a colony occurring in a pixel given the other colonies in the pattern) with two components: a first-order term (“trend”, the conditional intensity for an empty point pattern) posing different forms of association (categorical, linear, quadratic) with the covariates describing vegetation heterogeneity (patch type and distance to their borders), and a second term modeling pairwise interactions among colonies as a hybrid Strauss-Hard core point process (Baddeley et al. 2013), with the “hard” component acknowledging restrictions of our sampling grain (colonies can never occur <1 m apart) and the “Strauss” component allowing for positive (Strauss interaction parameter ɣ > 1) or negative (ɣ < 1) interactions up to a critical interaction radius of 6 m (determined by likelihood profiling; see DOI:10.5281/zenodo.15521695). Gibbs models were fitted to data by pseudolikelihood (composite likelihood) methods (Baddeley et al. 2016) and they were compared with the Akaike Information Criterion (AIC, which penalizes model complexity), with ΔAIC > 2 suggesting a significant change favoring the model with the lowest AIC value.

## Results

### Spatiotemporal trends in colony density

Densities of *Pheidole bergi* colonies were similar within the algarrobal and on dirt roads crossing it when sampled in the same season (2019). However, strong density changes were observed between seasons, with a >50% reduction in both permanent grids between the wet 2001 and the dry 2019, and a further >40% reduction in both dirt roads along the drought from 2019 to 2025 (Fig. 1). In 2021, colony densities were higher in every dirt road segment crossing the algarrobal than in those crossing the sand dunes or the grazed algarrobal (Fig. 1).

**Fig. 1.**
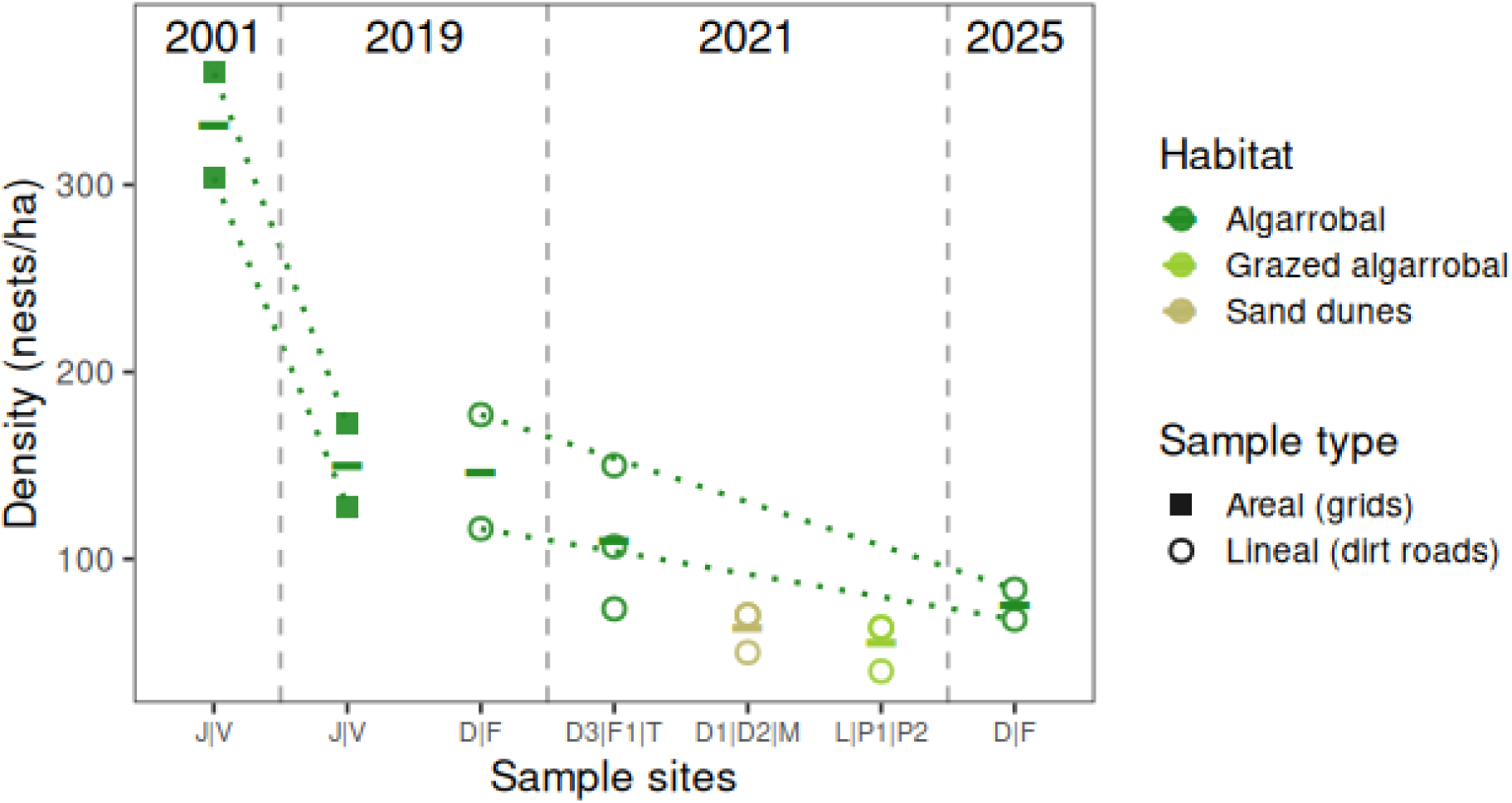
Density of colonies of *Pheidole bergi* in permanent grids on the algarrobal (squares) and on dirt roads crossing different habitats (circles) of the Ñacuñán Reserve and an adjacent cattle ranch in 2001, 2019, 2021 and 2025. Horizontal bars indicate the mean value for each group of samples; dotted lines join censuses of the same permanent grid and the same dirt road. Note that in the two last groups for 2021 there are overlapping circles for pairs of samples with the same estimated densities.

### Colony distribution along dirt roads (macro scale)

We found strong evidence that the intensity of the linear spatial process was significantly heterogeneous in both 2019 and 2025 along the dirt road crossing the algarrobal that had more colonies of *P. bergi* (D; 186 and 88 colonies / 1500 m in 2019 and 2025, respectively), where colonies appeared clumped on scales at least up to 150 m (Fig. 2). In contrast, no significant heterogeneity was detected along road F, although the degree of aggregation at most distances was higher in 2025, with less colonies (77 colonies / 1500 m), than in 2019 (122 colonies / 1500 m; Fig. 2). Two of the three shorter segments of roads crossing the algarrobal sampled in 2021 also showed strong evidence of colony aggregation at distances of at least 60–100 m (Fig. 3). Dirt roads crossing the sand dunes and the grazed algarrobal had around half the number of active colonies, more homogeneously distributed (Fig. 1); only one of the six segments (M, crossing the sand dunes) showed heterogeneous intensity with all but one colony concentrated in half of the transect (Fig. 3).

**Fig 2.**
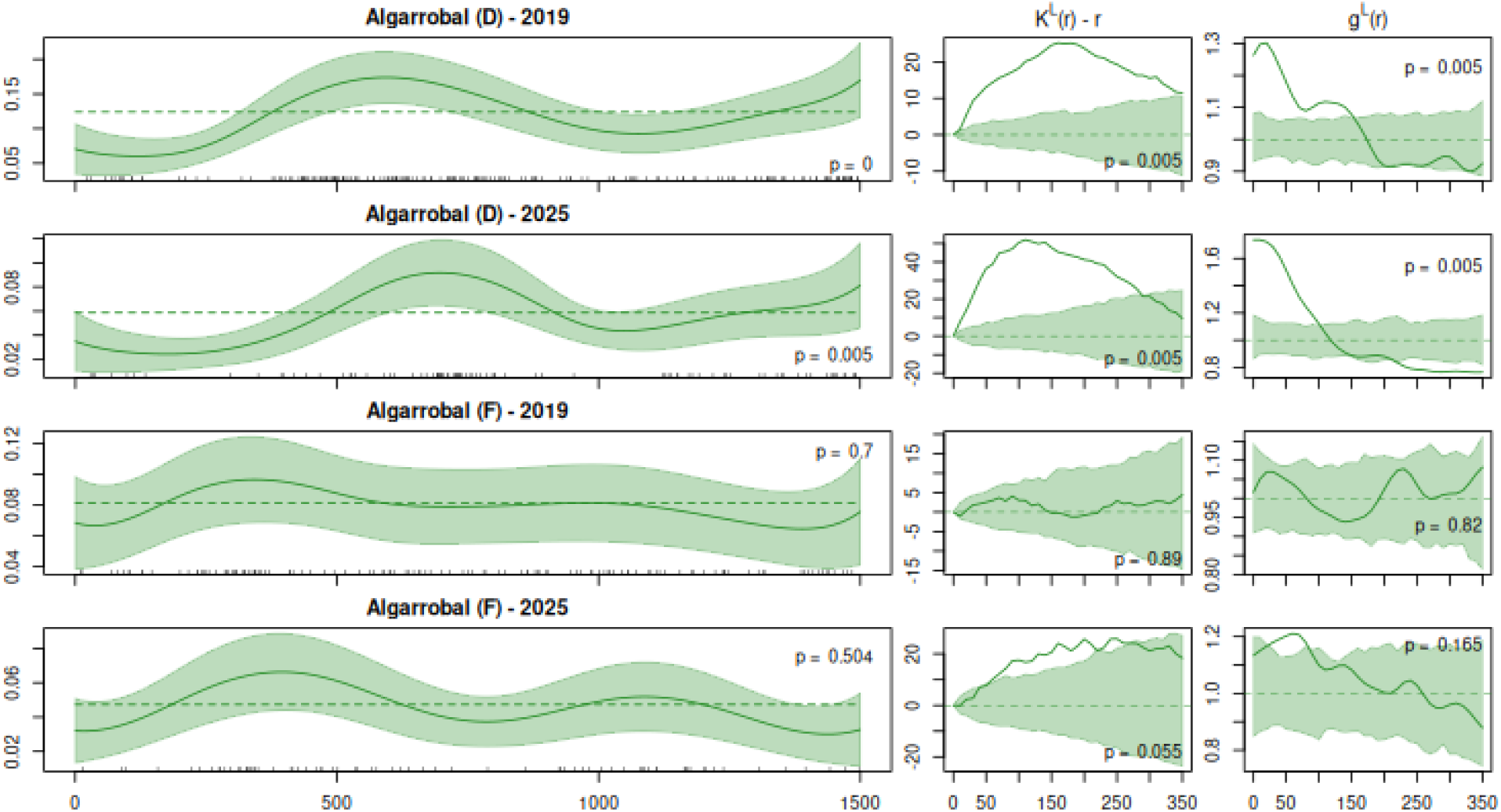
Spatial distributions of *Pheidole bergi* colonies along 1500-m segments of two dirt roads (D, F) crossing the algarrobal of the Ñacuñán Reserve in 2019 and 2025. Left: colony positions (black ticks over x axis) and estimated intensity of the corresponding Poisson linear point pattern process when assumed homogeneous (dashed line) or variable along each segment (estimated by non-parametric gaussian kernel smoothing, continuous line, with shading for its 95% confidence interval); p-value corresponds to a cumulative distribution function test for homogeneous intensity. Right: spatial correlation of the colonies according to Ripley’s K (centered) and pair correlation (*g*) functions (line: observed; shade: 95% critical pointwise envelope assuming homogeneous Poisson processes) of the distance among colonies (*r*, in m); p-value corresponds to a global Maximum Absolute Deviation test that the process is Poisson (= no spatial correlation).

**Fig 3.**
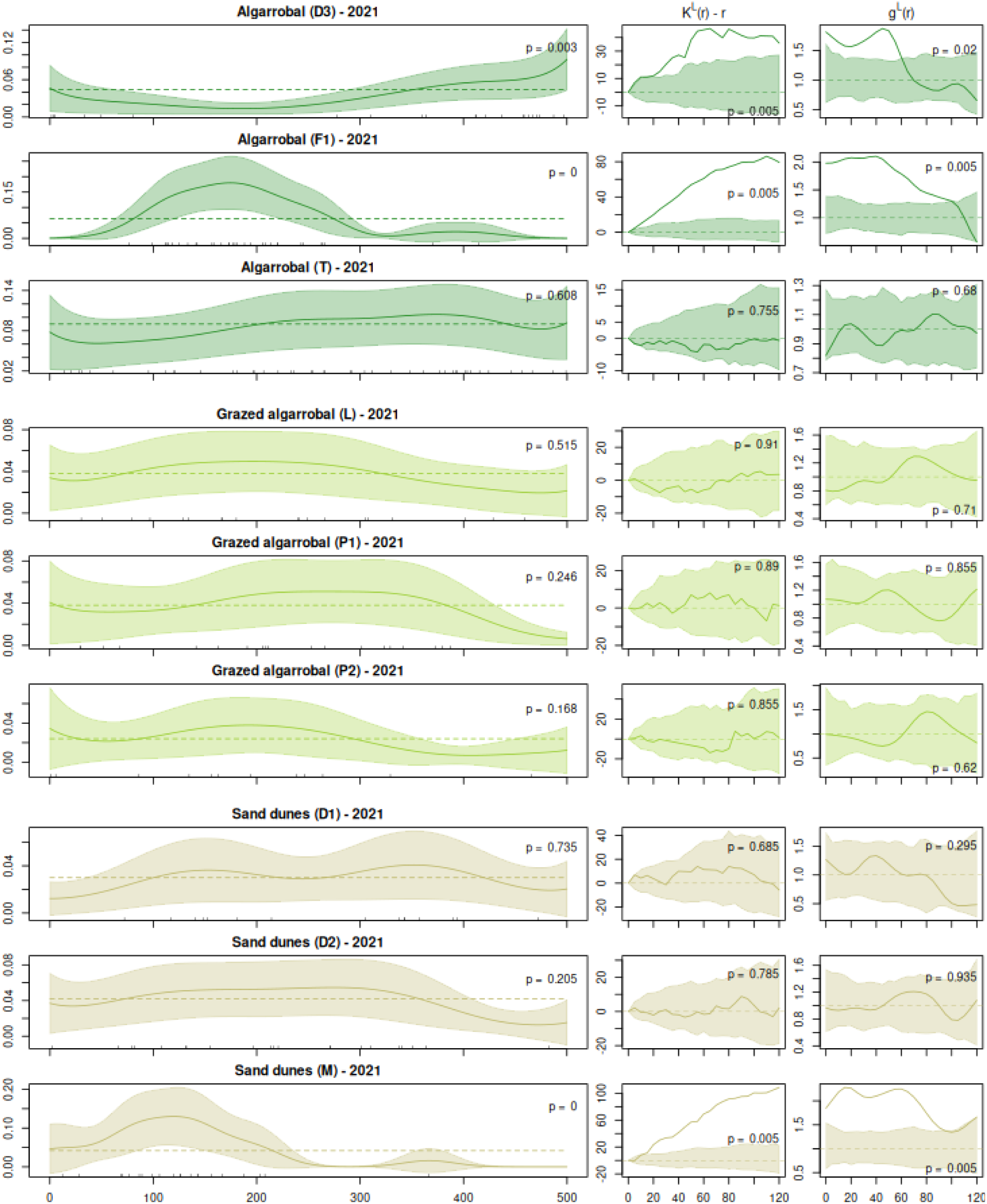
Spatial distributions of *Pheidole bergi* colonies along 500-m segments of dirt roads crossing the algarrobal of the Ñacuñán Reserve (D3, F1, T), a grazed algarrobal in an adjacent cattle ranch (L, P1, P2), and sand dunes within the reserve (D1, D2, M) in 2021. See Fig. 2 for references on subfigures, symbols and statistical values.

All road segments with significant colony aggregation (D in 2019 and 2025; D3, F1 and M in 2021) were well described by Poisson models with variable intensity along the roads (shown in Figs. 2 and 3) that assume no residual autocorrelation (i.e., distance-dependent pairwise interactions) among colonies (all tests based on inhomogeneous K and inhomogeneous pair correlation functions: p > 0.1; see DOI:10.5281/zenodo.15521695).

### Colony distribution in the algarrobal (micro scale)

Although colony densities consistently differed between the permanent grids J and V (Fig. 1), both showed clumped distributions of nests in the wet year 2001: there were more colonies than expected for a CSR process (*g* ≥ 1) at all explored distances resulting in significative aggregation at all scales bigger than 4–5 m (pointwise L above the CSR envelope; Fig. 4). In 2019, with less than half of the colonies (Fig. 1), the spatial pattern in V shifted to a repulsive or uniform pattern up to 6–7 m, while spatial autocorrelation in J became more homogeneous (i.e., not different than expected under CSR; Fig. 4).

**Fig 4.**
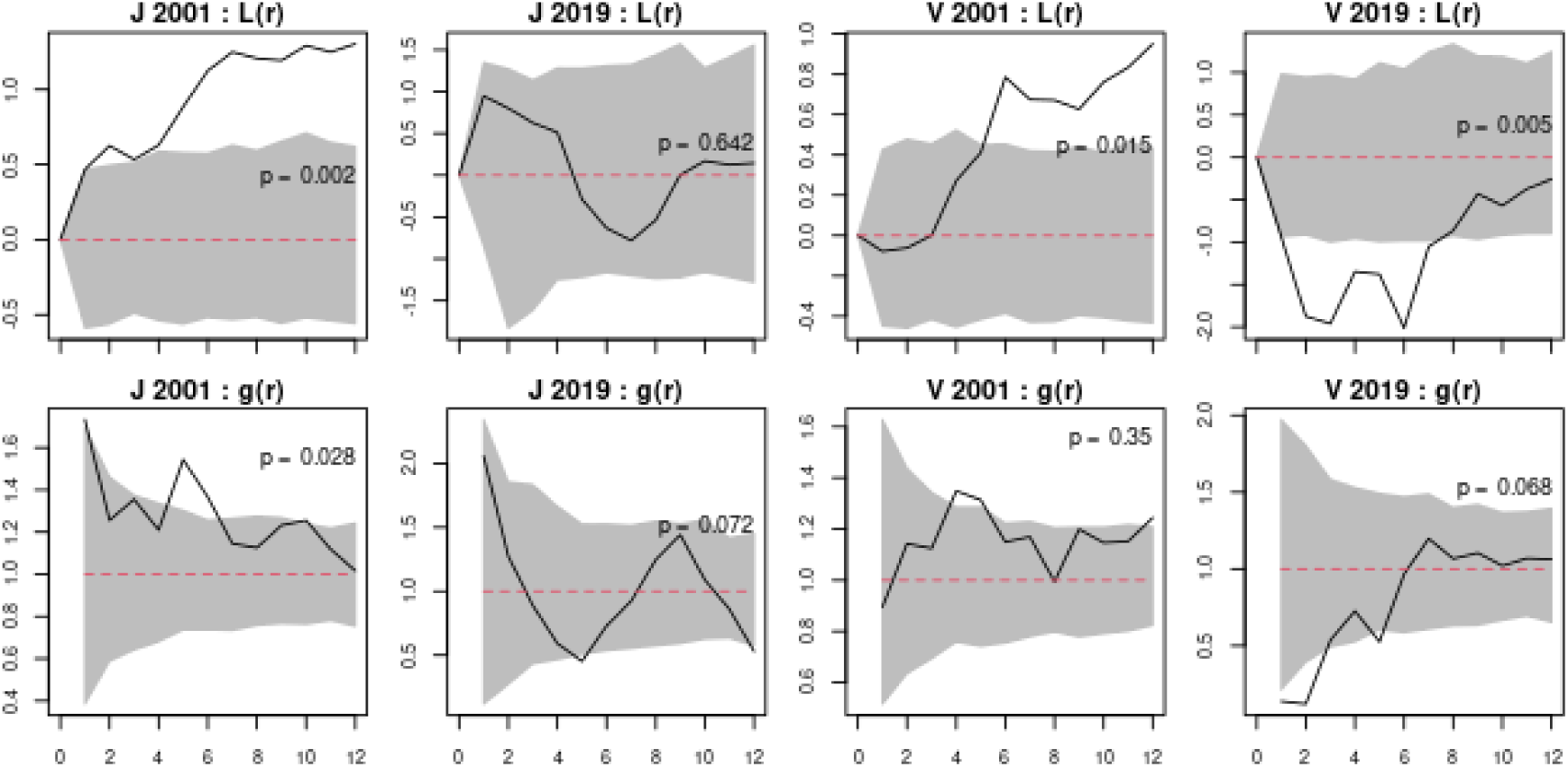
Spatial correlation of the *Pheidole bergi* colonies in two permanent 50×50 m grids (J, V) in the algarrobal of the Ñacuñán Reserve in 2001 and 2019, according to the two dimensional Ripley’s L function (above) and the pair correlation function (*g*, below) of the distance among colonies (*r*, in m). Red dashed lines are the expected values for a CSR process, black lines the observed values and shaded areas the 95% critical envelopes for pointwise tests of a CSR process with the same intensity; p-value corresponds to a global Maximum Absolute Deviation test that the spatial process is CSR (i.e., with homogeneous intensity and no spatial interactions among colonies).

Vegetation in the two permanent grids was generally similar, although tall woody cover (>1 m high) was slightly higher in J (2001: 61%; 2019: 69%) than in V (2019: 58%) and grid V contained a larger vegetation patch (allowing for greater “negative” distances to inter-patch borders; Fig. 5). In grid J, pixels with *P. bergi* colonies showed a non-random distribution of distances to the inter-patch borders, being more associated with open patches; in grid V similar tendencies were observed but within the expectations of a random process (Fig. 5).

**Fig 5.**
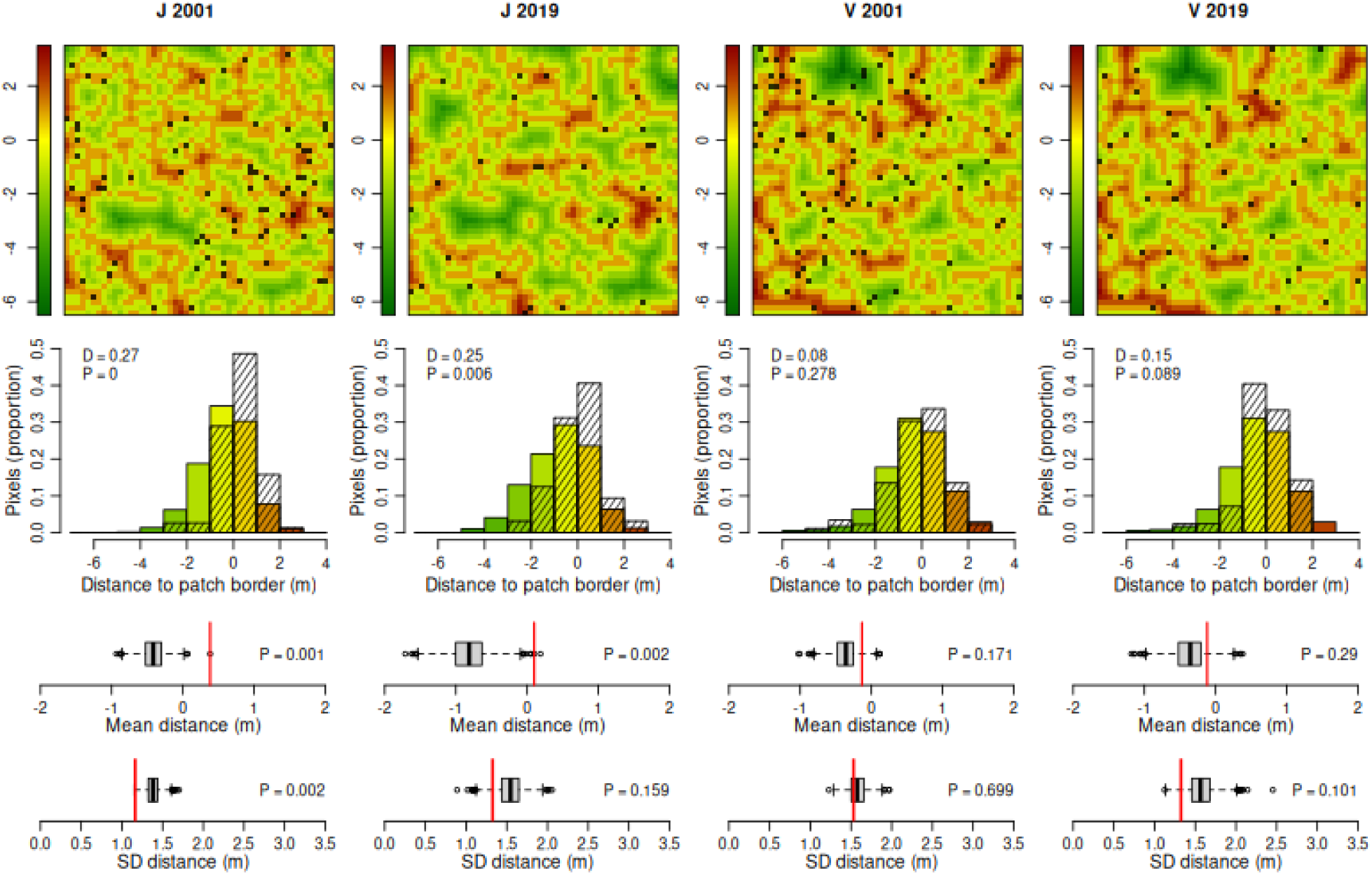
Top: distance to the nearest inter-patch border (negative values for pixels within woody vegetation patches, positive values for pixels located in open patches) of each 1 m^2^ pixel in two permanent 50×50 m grids (J, V) in the algarrobal of the Ñacuñán Reserve in 2001 and 2019; black pixels are those with *Pheidole bergi* colonies. Second row: frequency distribution of all pixels (n = 2500, coloured bars) in each distance category and of those pixels with colonies (hatched bars), with statistics corresponding to the Kolmogorov-Smirnov goodness-of-fit tests. Below: statistics of position (mean, third row) and dispersion (SD, fourth row) of the observed distribution of distances for pixels with colonies (red vertical line) and those obtained from 1999 random samples (of the same size) from the available distances (box and whiskers); the empirical p-value tests CSR as null model.

All Poisson point process models incorporating heterogeneous intensity based on the location and type of vegetation fitted data better than models assuming homogeneous intensities for each grid (CSR; Table 1). Among the tested associations with vegetation, models with intensity potentially varying both with the distance to the nearest inter-patch border and the type of patch (quadratic relationship with distance to inter-patch border: DIST^2^; different linear slopes for each type of patch: BRDHET) provided better fits than those models considering only one of those relationships with vegetation (only patch type: HET; single slope with distance: BRD). All models including distance-related interactions between colonies (Strauss-Hard models), either positive (attractive) or negative (repulsive), fitted better than their corresponding Poisson models with the same structure but no interactions (e.g., AIC_HOM_ < AIC_CSR_, AIC_HET+_ < AIC_HET_, AIC_BRD+_ < AIC_BRD_; Table 1). Importantly, the spatial effect of vegetation patches on *P. bergi* colonies was still relevant with inter-colony interactions allowed (e.g., AIC_HET+_ < AIC_HOM_; AIC_BRD+_ < AIC_HOM_), preserving their relative likelihoods within the set (AIC_BRDHET+_ < AIC ^2^ < AIC_HET+_ < AIC_BRD+_; Table 1).

**Table 1.** Structure and parameters for Poisson and Gibbs point process models fitted to the distributions of the *Pheidole bergi* colonies in two permanent 50×50 m grids in the algarrobal of the Ñacuñán Reserve in 2001 and 2019, considered replicates of the same spatial process. CSR is a Complete Spatial Random Model (= homogeneous intensity for each year, assuming independence between points) while HOM+ is the corresponding Gibbs model with a homogeneous first order term and interaction intensity varying in time. For the rest of the fitted models, those allowing for inter-point interactions are identified with a “+” following the name of the corresponding Poisson model with the same structure for the first-order term. In the Intensities/Trends column, “m” is the value of distance (in m) to the nearest inter-patch border (with negative values for distances toward the interior of the woody vegetation patches). The model with the lowest Akaike’s Information Criterion (AIC) value in the set is shown in bold.

| Model | Structure | Interactions | Intensities/Trends | AIC |
| --- | --- | --- | --- | --- |
| CSR | year | — | $\beta_{2001}: 0.033$<br>$\beta_{2019}: 0.015$ | 2231.25 |
| HOM+ | year | $\gamma_{2001}: 1.147$<br>$\gamma_{2019}: 0.53$ | $\beta_{2001}: 0.019$<br>$\beta_{2019}: 0.033$ | 2166.97 |
| HET | year + veg/open | — | 2001: $\beta_{\text{veg}}: 0.024$ / $\beta_{\text{open}}: 0.046$<br>2019: $\beta_{\text{veg}}: 0.011$ / $\beta_{\text{open}}: 0.021$ | 2209.3 |
| HET+ | year + veg/open | $\gamma_{2001}: 1.138$<br>$\gamma_{2019}: 0.503$ | 2001: $\beta_{\text{veg}}: 0.014$ / $\beta_{\text{open}}: 0.027$<br>2019: $\beta_{\text{veg}}: 0.027$ / $\beta_{\text{open}}: 0.053$ | 2142.3 |
| BRD | year + abs(dist) | — | 2001: $0.061 \times 0.622^{\text{m}}$<br>2019: $0.029 \times 0.622^{\text{m}}$ | 2216.34 |
| BRD+ | year + abs(dist) | $\gamma_{2001}: 1.149$<br>$\gamma_{2019}: 0.49$ | 2001: $0.038 \times 0.571^{\text{m}}$<br>2019: $0.082 \times 0.571^{\text{m}}$ | 2144.63 |
| BRDHET | year + abs(dist) *<br>veg/open | — | 2001: $\beta_{\text{veg}}: 0.052 \times 0.583^{\text{m}}$ / $\beta_{\text{open}}: 0.052 \times 0.891^{\text{m}}$<br>2019: $\beta_{\text{veg}}: 0.025 \times 0.583^{\text{m}}$ / $\beta_{\text{open}}: 0.025 \times 0.891^{\text{m}}$ | 2200.03 |
| **BRDHET+** | year + abs(dist) *<br>veg/open | $\gamma_{2001}: 1.14$<br>$\gamma_{2019}: 0.473$ | 2001: $\beta_{\text{veg}}: 0.033 \times 0.537^{\text{m}}$ / $\beta_{\text{open}}: 0.033 \times 0.839^{\text{m}}$<br>2019: $\beta_{\text{veg}}: 0.073 \times 0.537^{\text{m}}$ / $\beta_{\text{open}}: 0.073 \times 0.839^{\text{m}}$ | 2126.89 |
| DIST<sup>2</sup> | year + dist<sup>2</sup> | — | 2001: $0.039 \times 1.238^{\text{m}} \times 0.936^{\text{m}^2}$<br>2019: $0.018 \times 1.238^{\text{m}} \times 0.936^{\text{m}^2}$ | 2202.6 |
| DIST<sup>2</sup>+ | year + dist<sup>2</sup> | $\gamma_{2001}: 1.139$<br>$\gamma_{2019}: 0.474$ | 2001: $0.023 \times 1.251^{\text{m}} \times 0.918^{\text{m}^2}$<br>2019: $0.052 \times 1.251^{\text{m}} \times 0.918^{\text{m}^2}$ | 2130.04 |

According to the best-fitting model (BRDHET+; Table 1), the base probability of an active colony was highest at inter-patch borders, declining 56% faster (linearly in the scale of the model) into the vegetated patch than towards the open patch, strongly favoring the presence of colonies in the latter (Fig. 6). The interaction between pairs of colonies (1–6 m apart) was slightly positive in 2001, favoring nest aggregation, but became strongly negative in 2019, favoring more regular patterns (particularly in V; Fig. 5). Gibbs point process models with alternative structures showed reduced likelihoods (higher AIC values) than BRDHET+, such as a model with a time-invariant inhomogeneous first order trend and differences between years only depending on intercolony interactions (∼abs(dist) * veg/open + Strauss-Hard_Year_, AIC = 2137.8), or with inhomogeneous first-order trend but a constant parameter for interactions (∼Year + abs(dist) * veg/open + Strauss-Hard, AIC = 2183.7). A more complex model with temporal changes in the vegetation effect on the first order trend was not significantly better than with the single, permanent relationship modeled in BRDHET+ (∼Year * abs(dist) * veg/open + Strauss-Hard_Year_, AIC = 2127.78).

**Fig 6.**
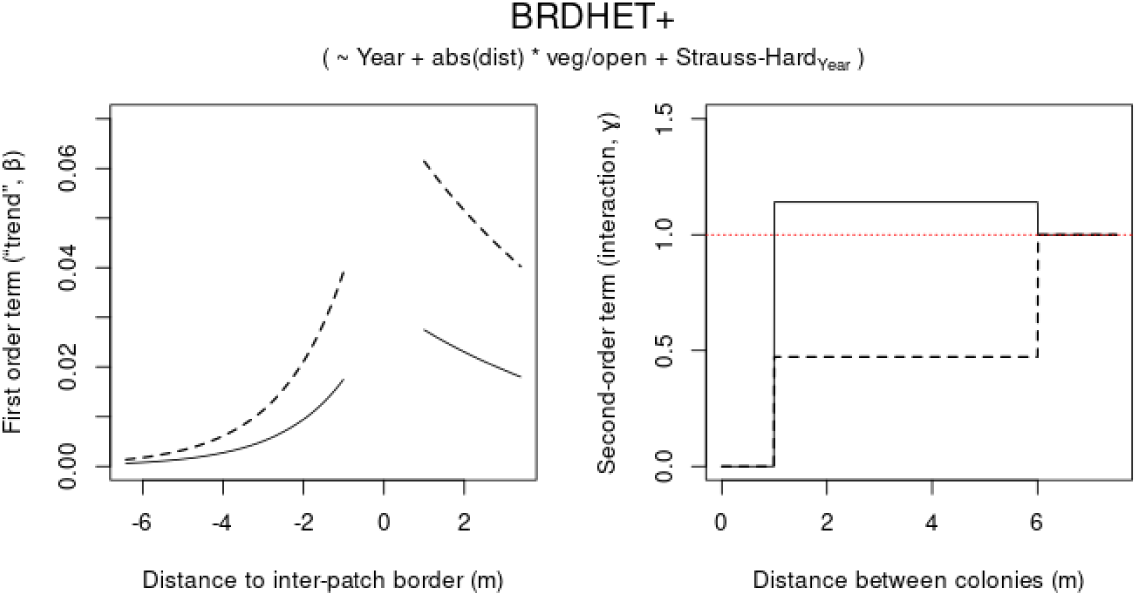
Selected Gibbs point process model (BRDHET+, Table 1) to explain the spatial distribution of *Pheidole bergi* colonies in the algarrobal of the Ñacuñán Reserve. Its first-order term (left) is the conditional intensity of the spatial pattern of the colonies that depends on time (2001: continuous line; 2019: dashed line) and distance to the nearest inter-patch border (negative values for pixels within woody vegetation patches). Its second-order term (right) is a Strauss-Hard model with interactions between active colonies up to 6 m (ɣ > 1: positive or attractive; ɣ < 1: negative or repulsive) also depending on time.

## Discussion

*Pheidole bergi* is not only very abundant in the Ñacuñán Reserve, as we had some previous evidence (Pirk et al. 2009; Miretti et al. 2025), but with a colony density far exceeding previous records for the species and for several other arid and semiarid ant species in South America. The density we recorded in its preferred habitat, the algarrobal (67–360 colonies/ha), was much higher than reported for the same species in the Caldenal woodland (8 colonies/ha; Tizón et al. 2010) and for all other ant species with available data in the central Monte (*Pogonomyrmex mendozanus*, *P. inermis, P. rastratus*, *Acromyrmex striatus*, *A. lobicornis*; Nobua Behrmann 2014; Pirk et al. 2004; Pol et al. 2017). Moreover, the colony densities we recorded at its peak during the wet period including 2001 (304–360 colonies/ha) compares to the highest values reported for entire seed-harvesting ant assemblages in some North American deserts: 340–520 colonies/ha in the Chihuahuan desert (Schumacher and Whitford 1976) and 61–463 colonies/ha in the Mojave and Great Basin deserts (Bernstein and Gobbel 1979). With its high densities, omnivorous diet, aggresive behaviour and environmental tolerance, *P. bergi* may play a relevant ecological role within the ant assemblage in South American arid and semi-arid ecosystems as other species of the genus have been recognized elsewhere (e.g., “an effective ecosystem engineer” in tropical agroecosystems: Shukla et al. 2013).

At the landscape scale, *P. bergi* showed a clear spatial preference between habitats and within its preferred habitat, with colony densities in the algarrobal doubling those in dirt roads crossing sand dunes or grazed algarrobal and with strong and persistent variations in colony density along most road segments crossing the algarrobal. The lower density of colonies in the grazed woodland may be associated with its reduced cover of the lowest vegetation strata (Milesi et al. 2002; Andrade 2016) or to the negative effect of cattle grazing on arthropod abundance, the main food source of *P. bergi* (Pirk et al. 2009), as commonly observed in arid ecosystems (e.g., Cheli and Martínez 2017; Pol et al. 2017; Martínez et al. 2021). The sandy and permeable dune soils, with low water retention, seemed to also limit the establishment and survival of *P. bergi*, as another case of soil texture critically affecting the abundance and distribution of ant species in arid and semiarid lands (e.g., Kirkham and Fisser 1972; Johnson 1992, 2000; Boulton et al. 2005). Compared to the preferred habitat, the protected algarrobal, both extensive limiting factors (anthropic: domestic grazing, natural: geomorphology) had homogenously reduced mean densities rather than the expected increased clumpiness resulting from (fewer) successful colonies occupying only a few suitable patches. The low number of colonies may also hinder the detection of weak non-random spatial patterns (lower power to test CSR as null hypothesis). Only one segment over sand dunes was the exception, with a radically clumped pattern of all but one colony in half of the transect. In contrast, most segments crossing the protected algarrobal showed aggregated patterns at scales of tens to a hundred meters, maybe as response to subtler variations in environmental constraints such as soil or vegetation characteristics. The segregated distribution of colonies of two granivorous ant species has been already associated with variations in soil texture and microtopography along these same 1500-m road segments within the algarrobal of Ñacuñán: *Pogonomyrmex inermis* associated with lower sectors and soils with higher clay content, and *P. mendozanus* with looser, coarser-textured soils in relative highs (Pol 2008; Milesi et al. MS).

A particularly striking result of our study was the marked decline in the abundance of *P. bergi* over the duration of our study. Colony density at the same sites halved between the wet 2001 and the dry 2019, but also decreased by at least another 40% during the multi-year drought between 2019 and 2025. Although these fluctuations could be part of the natural population dynamics of *P. bergi*, the magnitude and persistence of this decrease suggest a true population drop, mirroring recent reports of a global decline in insect populations (Sánchez-Bayo and Wyckhuys 2019; Møller 2020). This strong decline could result from reduced soil moisture and lower herbaceous plant cover associated with the persistent drought in this area since 2018 (Miretti et al. 2024; Vullo et al. 2024; Milesi et al, unpublished data), and their probable negative effect on the arthropods on which *P. bergi* feeds. Negative responses to drought and anthropogenic disturbances have been observed in desert ants driven by factors such as low production of reproductive individuals, insufficient rainfall to trigger mating flights, higher desiccation risk, lower foundress survival, decreasing foraging activity under low food availability, and increased colony mortality (Sanders and Gordon 2004; Gibb et al. 2019; Parr and Bishop 2022; Sundaram et al. 2022). contrary to our expectation, the decline between 2019 and 2025 within the preferred habitat of *P. bergi* appeared inversely density-dependent, as in an Allee effect: the sectors of higher and lower density along the roads persisted, but the proportional reduction in the number of colonies was (slightly) higher in the latter, resulting in a (weak) increase in the degree of aggregation at the macro scale (see Fig. 2). Probably the suboptimal sectors (already with lower nest densities) were no longer able to sustain active *P. bergi* colonies after accumulating the negative effects of a multi-year drought (i.e., the pattern we expected for harsher habitats). In no scenario distance-related interspecific interactions seemed necessary to explain the resulting patterns at this scale.

In contrast with the extensive impact of domestic grazing, the intensive anthropic perturbation of dirt road construction, management and use (although with very low traffic), which results in striking microhabitat differences with surrounding environments (e.g., plant cover and composition, temperature, soil compaction), do not seem to significantly impact the establishment and survival of *P. bergi* colonies. Density in the disturbed areas was not higher either as reported for several desert ants (e.g., Farji-Brener 1996; Terranella et al. 1999; Pirk et al. 2004; Tizón et al. 2010; Uhey et al. 2021). This tolerance can be at least partially explained by its microhabitat selection within the algarrobal. At this scale, *P. bergi* consistently avoided the interior of woody vegetated patches (shrubs and trees >1 m tall), with more active colonies near their borders and in the surrounding open patches. Many desert ants select open nesting sites too (e.g., Carlson and Gentry 1973; Bernstein and Gobbel 1979; Bestelmeyer and Schooley 1999; Nicolai 2005). Factors such as lower soil temperature reducing foraging activity and food intake (Nobua Behrmann et al. 2017), slower egg development (Bar et al. 2022), or lower efficiency detecting and transporting food in litter accumulating under canopy (Bin Kalban et al. 2024) have been proposed for other species and may explain why *P. bergi* avoids dense vegetation. Borders of woody patches in the Monte desert may offer particularly suitable nesting and foraging conditions because the sparse external branches and inverted conical shape of *Larrea divaricata*, the shrub that dominate most of them, result in microclimatic conditions similar to those of small open patches. This preference for borders of vegetation and open patches nearby may explain why relatively narrow dirt roads (6–10 m wide), cleared and compacted but not much bigger than open patches within the algarrobal, still offer suitable nesting sites for *P. bergi*.

Once that “exogenous” or “induced” spatial dependence (*sensu* Fortin and Dale 2005) within the algarrobal was accounted for, significative inter-colony interactions became evident, and they shifted over time with changes in environmental conditions and population densities (we are not be able to separate those confounded factors, which are expected to be highly correlated in these extreme ecosystems anyway). In 2001, with rainfall above long-term average and colony density at its peak, colonies were more aggregated than accounted for by vegetation (weak inter-colony attraction), but colony distribution shifted towards random or uniform patterns (and intercolony influence to strongly repulsive) by 2019, during a multi-year drought, when densities halved. This is the opposite of the expected positive correlation between colony density and intercolony repulsion that would result (*via* intraspecific competition) in regular spatial patterns (Schooley and Wiens 2003). If the wet period before 2001 was optimal for reproduction of *P. bergi*, new young colonies with few workers and small foraging territories, probably aggregated through restricted dispersal of new founding queens from natal nests or mating sites (Rissing and Pollock 1989; Cole and Wiernasz 2002), may have coexisted during a (relatively short) period of relaxed competition and abundant resources. As conditions worsened and resources declined, intraspecific competition likely favoured larger, well-established colonies, leading to density-dependent thinning and increased inter-colony spacing up to 2019 (Hölldobler and Wilson 1990; Wiernasz and Cole 1995). The proposed mechanism, then, is that stricter environmental conditions resulted in both reduced densities and increased intraspecific competition, especially in the most densely occupied areas (density-dependent thinning). It should be noted, however, that the two components of these Gibbs spatial models (first-order: trend; second-order: inter-colony interactions) to account for the distribution of *P. bergi* colonies within the algarrobal are partially confounded, as both operate at a similar spatial scale (maximum distance to an inter-patch border: 6.4 m; maximum distance for interactions: 6 m). In fact, allowing for interactions to vary with time inverted the estimated intercepts for the first-order term compared to those in the Poisson model that assume no distance-related interactions (BRDHET: β_2001_ > β_2019_; BRDHET+: β_2001_ < β_2019_; see Table 1 and Fig. 6). Maybe similar analyses in areas with bigger vegetation patches (particularly the preferred open patches) would allow for a clearer separation of those two components.

In summary, *P. bergi* is a highly abundant and potentially ecologically significant ant species in the central Monte desert. This ant species behaves as a diet and habitat generalist, flexible enough to establish colonies in different habitats and microhabitat conditions although with distribution and population dynamics clearly shaped by environmental conditions such as vegetation cover, soil texture or resource availability. Colony abundance varied across habitats, with lower densities in livestock-grazed ranches and sand dunes and high but spatially variable colony density in protected open woodlands and dirt roads that cross them. Within the algarrobal, *P. bergi* selects inter-patch borders and open patches, avoiding the interior of woody patches. The strong decline in density along the last decades with associated changes in spatial distribution suggest that both direct and indirect effects of environmental variability, including prolonged drought, play a key role in the establishment and survival of its colonies. Furthermore, the correlated inverse density-dependent thinning at the habitat scale and shifts in relevance of distance-dependent inter-colony interactions (or density-dependent interspecific competition) at smaller scales highlight the complex interplay between environmental constraints and intraspecific dynamics in the population ecology of *P. bergi*.

## Author contributions

**ALC**: Writing– original draft; writing– review and editing; formal analysis; investigation. **FAM**: Conceptualization; writing– original draft; writing– review and editing; formal analysis; investigation. **RGP**: Conceptualization; writing– review and editing; investigation; funding acquisition. **LV**: Writing– review and editing; investigation. **MFM**: Writing– review and editing; investigation. **GIP**: Writing– review and editing; investigation. **JLC**: Conceptualization; writing– review and editing; investigation; funding acquisition.

## Acknowledgements

We thank the Ñacuñán park rangers Pablo Mastrángelo, Raúl Porras, Ricardo Abraham, Alejandro Traslaviña, Mariela Valenzuela and Andrés Castro for their kind hospitality and logistical assistance. Adrian Baddeley provided invaluable assistance for the best use of the *spatstat* family of packages. We thank Luis Calcaterra, Julieta Filloy and Roxana Josens for their helpful comments on an earlier version of this manuscript. Financial support was provided by Universidad de Buenos Aires (more recently through UBACYT 20020220100224BA), Consejo Nacional de Investigaciones Científicas y Técnicas (more recently through PIP 2718) and Agencia Nacional de Promoción Científica y Tecnológica (more recently through PICT 3217), all from Argentina. This is contribution number XXX of the Desert Community Ecology Research Team (Ecodes) of IADIZA Institute (CONICET) and FCEN (Universidad de Buenos Aires).

## References

Aguiar MR and Sala OE (1999) Patch structure, dynamics and implications for functioning of arid ecosystems. Trends in Ecology and Evolution 14:273–277

Andrade LE (2016) Reclutamiento de plantas herbáceas en el desierto del Monte central: los papeles de la granivoría, la herbivoría y la vegetación leñosa. Doctoral thesis, Universidad de Buenos Aires, Buenos Aires

Ang QW, Baddeley A and Nair G (2011) Geometrically corrected second order analysis of events on a linear network, with applications to ecology and criminology. Scandinavian Journal of Statistics 39:591–617

Baddeley A, Diggle PJ, Hardegen A, Lawrence T, Milne RK and Nair G (2014) On tests of spatial pattern based on simulation envelopes. Ecological Monographs 84:477–489

Baddeley A, Hardegen A, Lawrence T, Milne RK, Nair G and Rakshit S (2017) On two-stage Monte Carlo tests of composite hypotheses. Computational Statistics and Data Analysis 114:75–87

Baddeley A, Rubak E and Turner R (2016) Spatial point patterns. Methodology and applications with R. CRS Press, Florida

Baddeley A, Turner R, Mateu J, and Bevan A (2013) Hybrids of Gibbs Point Process Models and Their Implementation. Journal of Statistical Software 55:1–43

Bar A, Shalev L and Scharf I (2022). Egg-laying behavior of *Cataglyphis niger* ants is influenced more strongly by temperature than daylength. Biology 11:art17149

Barrientos G, Dettler A, Martínez E, Vazquez F, Ansa A, Santadino M, Ewens M, Craig E and Riquelme Virgala M (2024) Variación espacial y temporal del ensamble de Formicidae en distintos tipos de bosque de la ecorregión Chaco Seco, Argentina. Ecología Austral 34:25–35

Bernstein RA and Gobbel M (1979) Partitioning of space in communities of ants. Journal of Animal Ecology 48:931–942

Bertiller MB, Marone L, Baldi R and Ares JO (2009) Biological interactions at different spatial scales in the Monte desert of Argentina. Journal of Arid Environments 73:212–221

Bestelmeyer BT and Schooley RL (1999) The ants of the southern Sonoran desert: community structure and the role of trees. Biodiversity and Conservation 8:643–657

Bestelmeyer BT and Wiens JA (2001) Ant biodiversity in semiarid landscape mosaics: the consequences of grazing vs natural heterogeneity. Ecological Applications 11:1123–1140

Bin Kalban M, Al Hammadi A and Bartholomew A (2024) Vegetation reduces the foraging efficiency of desert ants *Cataglyphis urens*, and they prefer unvegetated microhabitats. Journal of Arid Environments 220:art105104

Bisigato AJ, Villagra PE, Ares JO and Rossi BE (2009) Vegetation heterogeneity in Monte Desert ecosystems: a multi-scale approach linking patterns and processes. Journal of Arid Environments 73:182–191

Blanco-Moreno JM, Westerman PR, Atanackovic V and Torra J (2014) The spatial distribution of nests of the harvester ant *Messor barbarus* in dryland cereals. Insectes Sociaux 61:145–152

Boulton AM, Davies KF and Ward PS (2005) Species richness, abundance, and composition of ground-dwelling ants in northern California grasslands: role of plants, soil, and grazing. Environmental Entomology 34:96–104

Bruch C (1916) Contribución al estudio de las hormigas de la provincia de San Luis. Revista del Museo de La Plata 23:291–357

Calcaterra LA, Cabrera SM, Cuezzo F, Jiménez Peréz I and Briano JA (2010) Habitat and grazing influence on terrestrial ants in subtropical grasslands and savannas of Argentina. Annals of the Entomological Society of America 103:635–646

Carlson DM and Gentry JB (1973) Effects of shading on the migratory behavior of the Florida harvester ant, *Pogonomyrmex badius*. Ecology 54:452–453

Cheli GH and Martínez FJ (2017) Artrópodos terrestres, su rol como indicadores ambientales. Pp. 98–117 in: Udrizar Sauthier DE, Pazos GE and Arias AM (eds) Reserva de Vida Silvestre San Pablo de Valdés 10 años: protegiendo el patrimonio natural y cultural de Península Valdés Patagonia argentina. Fundación Vida Silvestre Argentina and CONICET, Buenos Aires and Puerto Madryn

Claver S, Silnik SL and Campón FF (2014) Response of ants to grazing disturbance at the central Monte Desert of Argentina: community descriptors and functional group scheme. Journal of Arid Land 6:117–127

Cole BJ and Wiernasz DC (2002) Recruitment limitation and population density in the harvester ant, *Pogonomyrmex occidentalis*. Ecology 83:1433–1442

Collins SL, Belnap J, Grimm NB, Rudgers JA, Dahm CN, D’Odorico P, Litvak M, Natvig DO, Peters DC, Pockman WT, Sinsabaugh RL and Wolf BO (2014) A multiscale, hierarchical model of pulse dynamics in arid-land ecosystems. Annual Review of Ecology, Evolution and Systematics 45:397–419

Crist TO and Wiens JA (1996) The distribution of ant colonies in a semiarid landscape: implications for community and ecosystem processes. Oikos 76:301–311

Dale MRT and Fortin MJ (2005) Spatial analysis: a guide for ecologists. Cambridge University press, Cambridge

Farji-Brener AG (1996) Posibles vías de expansión de la hormiga cortadora de hojas *Acromyrmex lobicornis* hacia la Patagonia. Ecología Austral 6:144–150

Fortin MJ (2020) Spatial structure in population data. Pp. 299–313 in: Murray DL and Sandercock BK (eds) Population ecology in practice. John Wiley & Sons, Oxford

Fortin MJ and Dale M (2005) Spatial analysis: a guide for ecologists. Cambridge University Press, New York

Guénard B, Weiser M, Gomez K, Narula N and Economo EP (2017) The Global Ant Biodiversity Informatics (GABI) database: a synthesis of ant species geographic distributions. Myrmecological News 24:83–89

Gibb H, Grossman BF, Dickman CR, Decker D and Wardle G (2019) Long-term responses of desert ant assemblages to climate. Journal of Animal Ecology 88:1549–1563

Goosey HB, Smith JT, O’Neill KM and Naugle DE (2019) Ground-dwelling arthropod community response to livestock grazing: implications for avian conservation. Environmental Entomology 48:856–866

Hierro JL, Muiño WA, Farji-Brener A, Cock MC, and Pearson DE (2023) Species introduction shifts a trait’s function from mutualism to antagonism: elaiosomes in a myrmecochory cold spot. Oikos 2023:e09770

Hölldobler B and Wilson EO (1990) The ants. Harvard University Press, Cambridge

Janicki J, Narula N, Ziegler M, Guénard B and Economo EP (2016) Visualizing and interacting with large-volume biodiversity data using client-server web-mapping applications: the design and implementation of antmaps.org. Ecological Informatics 32:185–193

Johnson RA (1992) Soil texture as an influence on the distribution of the desert seed-harvester ants *Pogonomyrmex rugosus* and *Messor pergandei*. Oecologia 89:118–124

Johnson RA (1998) Foundress survival and brood production in the desert seed-harvester ants *Pogonomyrmex rugosus* and *P. barbatus* (Hymenoptera, Formicidae). Insectes Sociaux 45:255–266

Johnson RA (2000) Habitat segregation based on soil texture and body size in the seed-harvester ants *Pogonomyrmex rugosus* and *P. barbatus*. Ecological Entomology 25:403–412

Johnson RA (2001) Biogeography and community structure of North American seed-harvest ants. Annual Review of Entomology 46:1–29

Kirkham DR and Fisser HG (1972) Rangeland relations and harvester ants in northcentral Wyoming. Journal of Range Management 25:55–60

Kusnezov N (1951) El género *Pheidole* en la Argentina (Hymenoptera, Formicidae). Acta Zoológica Lilloana 12:5–88.

Levings SC and Traniello JFA (1981) Territoriality, nest dispersion and community structure in ants. Psyche 88:265–319

Lopez de Casenave J (2001) Estructura gremial y organización de un ensamble de aves del desierto del Monte. Doctoral thesis, Universidad de Buenos Aires, Buenos Aires

MacKay WP (1991) The role of ants and termites in desert communities. Pp. 113–150 in: Polis GA (ed) The ecology of desert communities. University of Arizona Press, Tucson

Marone L, Cueto VR, Milesi FA and Lopez de Casenave J (2004) Soil seed bank composition over desert microhabitats: patterns and plausible mechanisms. Canadian Journal of Botany 82:1809–1816

Marone L and Horno ME (1997) Seed reserves in the central Monte Desert, Argentina: implications for granivory. Journal of Arid Environments 36:661–670

Marone L and Pol RG (2021) Continuous grazing disrupts desert grass-soil seed bank composition under variable rainfall. Plant Ecology 222:247–259

Martínez FJ, Dellapé PM, Bisigato AJ, Zaffaroni FT and Cheli GH (2021) Shrub-dwelling arthropod assemblages respond differently to grazing disturbance in the southern Monte, Argentina. Journal of Arid Environments 118:art104384

Milesi FA and Lopez de Casenave J (2004) Unexpected relationships and valuable mistakes: non-myrmecochorous *Prosopis* dispersed by messy leafcutting ants in harvesting their seeds. Austral Ecology 29:558–567

Milesi FA, Lopez de Casenave J and Cueto VR (2008) Selection of foraging sites by desert granivorous birds: vegetation structure, seed availability, species specific foraging tactics, and spatial scale. Auk 125:473–484

Milesi FA, Lopez de Casenave J and Cueto VR (2019) Are all patches worth exploring? Foraging desert birds do not rely on environmental indicators of seed abundance at small scales. BMC Ecology 19:art25

Milesi FA, Marone L, Lopez de Casenave J, Cueto VR and Mezquida ET (2002) Gremios de manejo como indicadores de las condiciones del ambiente: un estudio de caso con aves y perturbaciones del hábitat en el Monte central, Argentina. Ecología Austral 12:149–161

Milesi FA, Pol RG, Pirk GI and Lopez de Casenave J (MS) Environmental associations and spatial segregation among coexisting Pogonomyrmex species in the central Monte desert

Miretti MF (2022) Respuestas de las hormigas granívoras al pastoreo en el desierto del Monte: patrones y mecanismos a distintas escalas. Doctoral thesis, Universidad de Buenos Aires, Buenos Aires

Miretti MF, Pol RG, Paris CI, Elizalde Capellino V, Sosa RA and Lopez de Casenave J (2025) Diversity and composition of seed-carrying ant assemblages in the Monte desert: a regional assessment of the effects of grazing. Journal of Insect Conservation 29:61

Miretti MF, Pol R, Vullo L, Cao AL, Marone L and Lopez de Casenave J (2024) Diet flexibility in three harvester ants (*Pogonomyrmex* spp.): effects of grazing and natural variations in the availability of seeds. Canadian Journal of Zoology 102:367–379

Møller AP (2020) Quantifying rapidly declining abundance of insects in Europe using a paired experimental design. Ecology and Evolution 10:2446–2451

Nash MS, Bradford DF, Franson SE, Neale AC, Whitford WG and Heggem DT (2004) Livestock grazing effects on ant communities in the eastern Mojave Desert, USA. Ecological Indicators 4:199–213

Nash MS, Whitford WG, Bradford DF, Franson SE, Neale AC and Heggem DT (2001) Ant communities and livestock grazing in the Great Basin, USA. Journal of Arid Environments 49:695–710

Nicolai NC (2005) Plant community dynamics governed by red harvester ant (*Pogonomyrmex barbatus*) activities and their role as drought refugia in a semi-arid savanna. Doctoral thesis, Texas A&M University, College Station

Nobua Behrmann BE (2014) Interacciones tróficas entre dos especies simpátricas de hormigas cortadoras y el ensamble de plantas en el Monte central. Doctoral thesis, Universidad de Buenos Aires, Buenos Aires

Nobua-Behrmann BE, Lopez de Casenave J, Milesi FA and Farji-Brener A (2017) Coexisting in harsh environments: temperature-based foraging patterns of two desert leafcutter ants (Hymenoptera: Formicidae: Attini). Myrmecological News 25:41–49

Noy-Meir I (1973) Desert ecosystems: environment and producers. Annual Review of Ecology and Systematics 4:25–51

Parr CL and Bishop TR (2022) The response of ants to climate change. Global Change Biology 28:3188–3205

Pirk GI, Di Pasquo F and Lopez de Casenave J (2009) Diet of two sympatric *Pheidole* spp. ants in the central Monte desert: implications for seed-granivore interactions. Insectes Sociaux 56:277–283

Pirk GI, Lopez de Casenave J and Pol R (2004) Asociación de las hormigas granívoras *Pogonomyrmex pronotalis*, *P. rastratus* y *P. inermis* con caminos en el Monte Central. Ecología Austral 14:65–76

Pol RG, Pirk GI and Marone L (2010) Grass seed production in the central Monte desert during successive wet and dry years. Plant Ecology 208:65–75

Pol RG (2008) Granivoría por hormigas del género *Pogonomyrmex* en el Monte central: respuestas funcionales a las variaciones en la disponibilidad de semillas. Doctoral thesis, Universidad Nacional de Cuyo, Mendoza

Pol RG, Sagario MC and Marone L (2014) Grazing impact on desert plants and soil seed banks: implications for seed-eating animals. Acta Oecologica 55:58–65

Pol RG, Vargas GA and Marone L (2017) Behavioural flexibility does not prevent numerical declines of harvester ants under intense livestock grazing. Ecological Entomology 42:283–293

R Core Team (2025) R: a language and environment for statistical computing. R Foundation for Statistical Computing, Vienna

Radnan GN and Eldridge DJ (2018) Ants respond more strongly to grazing than changes in shrub cover. Land Degradation and Development 29:907–915

Rissing SW and Pollock GB (1989) Behavioural ecology and community organization of desert seed-harvester ants. Journal of Arid Environments 17:167–173

Sánchez-Bayo F and Wyckhuys KAG (2019) Worldwide decline of the entomofauna: a review of its drivers. Biological Conservation 232:8–27

Sanders NJ and Gordon DM (2004) The interactive effects of climate, life history, and interspecific neighbours on mortality in a population of seed harvester ants. Ecological Entomology 29:632–637

Schooley RL and Wiens JA (2003) Spatial patterns, density dependence, and demography in the harvester ant, *Pogonomyrmex rugosus*, in semi-arid grasslands. Journal of Arid Environments 53:183–196

Schumacher A and Whitford WG (1976) Spatial and temporal variation in Chihuahuan Desert ant faunas. Southwestern Naturalist 21:1–8

Shukla RZ, Singh H, Rastogi N and Agarwal VM (2013) Impact of abundant *Pheidole* ant species on soil nutrients in relation to the food biology of the species. Applied Soil Ecology 71:15–23

Sundaram M, Steiner E and Gordon DM (2022) Rainfall, neighbors, and foraging: the dynamics of a population of red harvester ant colonies 1988–2019. Ecological Monographs 92:e1503

Terranella AC, Ganz L and Ebersole JJ (1999) Western harvester ants prefer nest sites near roads and trails. Southwestern Naturalist 44:382–384

Tizón FR, Peláez DV and Elía OR (2010) Efecto de los cortafuegos sobre el ensamble de hormigas (Hymenoptera, Formicidae) en una región semiárida, Argentina. Iheringia 100:216–221

Tongway DJ and Ludwig JA (2005). Heterogeneity in arid and semiarid lands. Pp. 189–205 in: Lovett GM, Turner MG, Jones CG and Weathers KC (eds) Ecosystem function in heterogeneous landscapes. Springer, New York.

Uhey DA, Cummins GC, Rotter MC, Lassiter LS and Whitham TG (2021) Hiking trails increase abundance of harvester ant nests at Clear Creek, Arizona. Southwestern Entomologist 46:403–411

Villagra PE, Defosse GE, del Valle HF, Tabeni S, Rostagno M, Cesca E and Abraham E (2009) Land use and disturbance effects on the dynamics of natural ecosystems of the Monte Desert: implications for their management. Journal of Arid Environments 73:202–211

Vullo L, Lopez de Casenave J, Miretti MF, Cao AL, Marone L and Pol RG (2024) Changes in seed abundance driven by overgrazing and rainfall variability trigger variations in the diet of a small harvester ant. Ecological Entomology 49:191–204

Warburg I and Steinberger Y (1997) On the spatial distribution of nests of the ants *Messor arenarius* and *Messor ebeninus*. Journal of Arid Environments 36:671–676

Whitford WG and Duval BD (2020) Ecology of desert systems. 2nd edition. Academic Press,

Wiernasz DC and Cole BJ (1995) Spatial distribution of *Pogonomyrmex occidentalis*: recruitment, mortality and overdispersion. Journal of Animal Ecology 64:519–527

Wild AL (2007) A catalogue of the ants of Paraguay (Hymenoptera: Formicidae). Zootaxa 1622:1–55

Wilson EO (2003) Pheidole in the New World: a dominant, hyperdiverse ant genus. Harvard University Press, Cambridge

